# Characterization of low copy number human angiotensin-converting enzyme 2 (hACE2)-transgenic mice as an improved model of SARS-CoV-2 infection

**DOI:** 10.1101/2023.08.31.555800

**Authors:** Christine M. Bradshaw, Teodora Georgieva, Trevor N. Tankersley, Tama Taylor-Doyle, Larry Johnson, Jennifer L. Uhrlaub, David Besselsen, Janko Ž. Nikolich

**Affiliations:** Department of Immunobiology, University of Arizona College of Medicine -Tucson, Tucson, AZ; Arizona Center on Aging, University of Arizona College of Medicine -Tucson, Tucson, AZ; Genetically engineered mouse model (GEMM) Core, University of Arizona, Tucson, AZ; Arizona Animal Care, University of Arizona, Tucson, AZ; the BIO5 Institute, University of Arizona, Tucson, AZ; Aegis Consortium for Pandemic-Free Future, University of Arizona Health Sciences, Tucson, AZ

## Abstract

Coronaviridae are significant human pathogens, as evidenced by several outbreaks of severe respiratory infections in the past 20 years and culminating with the COVID-19 pandemic. Mouse models of COVID-19 have included transgenic expression of the main SARS coronavirus entry receptor on human cells, human angiotensin-converting enzyme 2 (hACE2). However, the original hACE2-Tg mouse strain overexpresses many copies of the transgene, leading to neuropathology not representative of human infection. Aiming to improve physiological relevance, we generated two new lines of hACE2-Tg mice using the original transgene construct expressing hACE2 under the control of the keratin 18 promoter (K18-hACE2).

We show that relative to the original strain, which expressed 8 copies of the transgene (8-hACE2-Tg), lines 1 and 2 expressed 1 and 2 copies of the transgene (1-hACE-2-Tg and 2-hACE-2-Tg, respectively). Upon intranasal (i.n.) infection with 10^3^ plaque-forming units (pfu) SARS-CoV-2 WA-1/US, 8-hACE2-Tg mice succumbed to infection by d. 7. 2-hACE2-Tg mice exhibited 31% survival, with less viral replication in the lung and brain when compared to 8-hACE2-Tg mice. Furthermore, SARS-CoV-2 infection in 1-hACE2-Tg mice exhibited no mortality and had no detectable virus in the brain, although they did show clear virus replication in the lung. All three mouse strains analyzed showed SARS-CoV-2-related weight loss that tracked with the mortality rates. 1-hACE2-Tg mice mounted detectable primary and memory T effector cell and antibody responses. We conclude that these strains, particularly 1-hACE2-Tg mice, provide improved models to study hACE2-mediated viral infections.

## Introduction

The significance of coronaviruses to human health needs no highlighting in the wake of the COVID-19 pandemic, that has left nearly 7 million dead as of early July 2023, and countless more with severe and/or protracted illness worldwide. Relevant animal models of coronavirus infection are of importance to understand the pathogenesis and host-microbe interactions for these viruses. In the infectious diseases research, inbred mouse strains are most frequently used to elucidate relevant questions regarding virus spread, pathogenesis and, in particular, immunity and immunogenetics, where a plethora of tools facilitates illuminating research. However, mice are not susceptible to natural infection by many sarbecoviruses, including SARS-CoV-2 ancestral and alpha to delta variants, due to incompatibility of the viral Spike glycoprotein with the mouse homolog of the main host entry receptor human angiotensin converting enzyme 2 (hACE2)^1,2,3^. To circumvent this problem, transgenic mice were generated expressing hACE2 under the keratin 18 (K18) promoter which directs transgene expression to epithelial cells, including those of the respiratory tract^4^. This model was successfully used to achieve infections with SARS and SARS-CoV-2 viruses and was instrumental in testing therapeutic and vaccine approaches to mitigate COVID-19^5,6,7,8,9^. In this model, ancestral SARS-CoV-2 infection in K18-hACE2 mice results in robust viral replication, severe pulmonary pathology, and mortality^10^. However, due to ectopic overexpression of hACE2, SARS-CoV-2-infected K18-hACE2 mice (shown to carry 8 copies of hACE2, and therefore designated as 8-hACE2-Tg in the text) also develop severe neurological disease with abundant viral replication in the brain contributing to neuroinflammation^10,11^ [although some groups (S. Perlman, personal communication) find that with aging the virus is no longer detectable in the brain]. This encephalitic pathology contributes critically to pronounced infection-associated mortality^12^. The intense SARS-CoV-2 replication in the brain of the 8-hACE2-Tg mouse model is not paralleled in humans, where one study concluded that “The lack of detectable SARS-CoV-2 by ddPCR or significant histologic evidence of direct infection suggests that active encephalitis is not a feature of COVID-19”^13^. Indeed, while hACE2 is expressed in several regions of the brain^14,15,16^, and while several autopsy reports have demonstrated the presence of SARS-CoV-2 RNA in brain tissue of some persons, expression on neurons is low ^17,18^, direct evidence for neuronal infection is largely missing, and most mortalities are due to pulmonary, vascular and other non-neurological complications^13,19,20^. Furthermore, while there is a potential for lethal outcomes upon SARS-CoV-2 infection, >90% of the patients recover from infection with only mild to moderate disease. Therefore, a mouse model that better approximates the tissue tropism and pathogenesis observed in human infection would be of value.

One strategy used to generate a suitable mouse model has been targeted mutagenesis to adapt SARS-CoV-2 to bind murine ACE2. Using this strategy, Baric’s group has produced SARS-CoV-2 MA10, a mouse-adapted virus that does not infect the brain but can cause lethal respiratory disease if given at high-enough doses^21^. As this strain differs genetically from the human strains (particularly from the variants evolving from the ancestral strain, that in mice cause no or very mild disease), developing additional models that use human strains is necessary.

The number of transgene copies inserted into the genome of transgenic mice is an important variable in the biology of transgenic mouse models and is often the source of phenotypic variation between different lines. 8-hACE2-Tg mice contain 8 transgene copies which has been shown to result in excess expression of hACE2, leading to excessive infection and rampant viral replication. The goal of the present study was to generate and characterize K18-hACE2 transgenic mice that express fewer copies of the transgene. We present the analysis of two such lines, bearing one and two transgene copies. We found direct correlation between the hACE2 transgene dose and survival, weight loss, and viral burden in the lung and brain. This demonstrates that reducing the hACE2 transgene copy number in mice reduces and even eliminates the confounder of excessive neuroinvasion by SARS-CoV-2 and provides an optimized model to interrogate pathogenesis by, and immunity to, coronaviridae that use hACE-2 as their primary entry receptor.

## Results

### Generation of low copy hACE2 transgenic mice

Commercially available K18-hACE2 mice (B6.Cg-Tg(K18-ACE2)2Prlmn/J; called 8-hACE2-Tg in the text were reported to express 8-9 copies of the K18-hACE2 transgene^4^. Previous studies have demonstrated that these mice exhibit high viral titers in the lungs and brain and rapidly succumb to severe pulmonary and neurological complications, dying between days 7 and 10 post infection (p.i.)^22,23,24^. We hypothesized that reducing transgene expression should limit the magnitude of the infection and result in disease progression more akin to that seen in humans. To address this hypothesis, we obtained the same gene cassette (generously provided by Drs Paul McCray and Stanley Perlman, University of Iowa, Iowa City, IA) employed to generate the original 8-hACE2-Tg strain^4^. We generated two founder lines of transgenic mice expressing 1 copy and 2 copies of the transgene designated as 1 and 2-hACE2-Tg, respectively. (See the Materials and Methods section and **Figure 1 A**). We used RT-qPCR of the hACE2 transcripts in the brain and lung to show that, when compared to the transcript levels found in 8-hACE2-Tg transgenic mice (Jackson Laboratory Strain #:034860), the newly generated lines expressed significantly lower hACE2 transcript levels, corresponding to 1 and 2 copies of hACE2, respectively (RT-qPCR, **Figure 1 B, C**).

**Figure 1.**
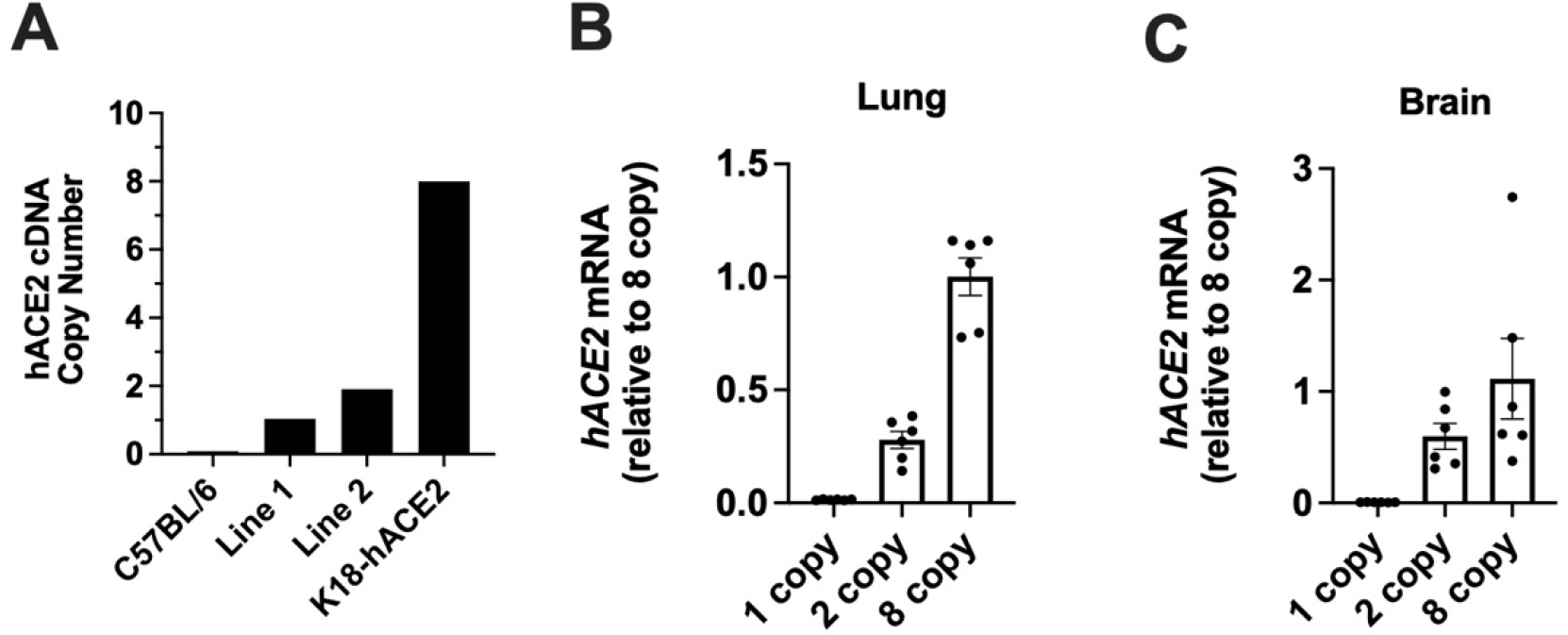
Generation of low copy hACE2 transgenic mice. (A) Quantitative PCR was used to determine hACE2 cDNA copy number in two transgenic founder lines and commercially available K18-hACE2 mice. (B) hACE2 mRNA expression in the lungs and (C) brains of 1-, 2-, and 8-hACE2-Tg mice. Results are mean ± SEM for 5 mice per group.

### SARS-CoV-2 pathogenesis depends on the hACE2 copy number in transgenic mice

To characterize SARS-CoV-2 infection in K18-hACE2 mice expressing fewer transgene copies, we intranasally (i.n.) infected mice expressing one, two, and 8 hACE2 transgene copies with 10^3^ PFU of the SARS-CoV-2 WA-1/US/2020 isolate, closely related to the ancestral Wuhan strain. Mice were then monitored daily for morbidity and changes in weight (**Figure 2 A, B**). While 8-hACE2-Tg mice exhibited ∼30% weight loss and 100% mortality by 11 days post infection (dpi), 1- or 2-hACE2-Tg mice experienced less severe disease. The 2-hACE2-Tg line exhibited a 31% survival rate and lost up to ∼12% of their starting weight, whereas the 1-hACE2-Tg line exhibited no mortality (**Figure 2 A, B**). Interestingly, 1-hACE2-Tg mice exhibited slight weight loss from day 6 p.i., which coincided with the time when the surviving 2-hACE2-Tg mice begin to regain their weight (**Fig 2B**). This may suggest a delay in disease onset in 1-hACE2-Tg mice. Body weight of the surviving 2-hACE2-Tg and all 2-hACE2-Tg mice returned to baseline by 45 d.p.i., suggesting a complete recovery after viral clearance in surviving mice (**Fig. 2B**).

**Figure 2.**
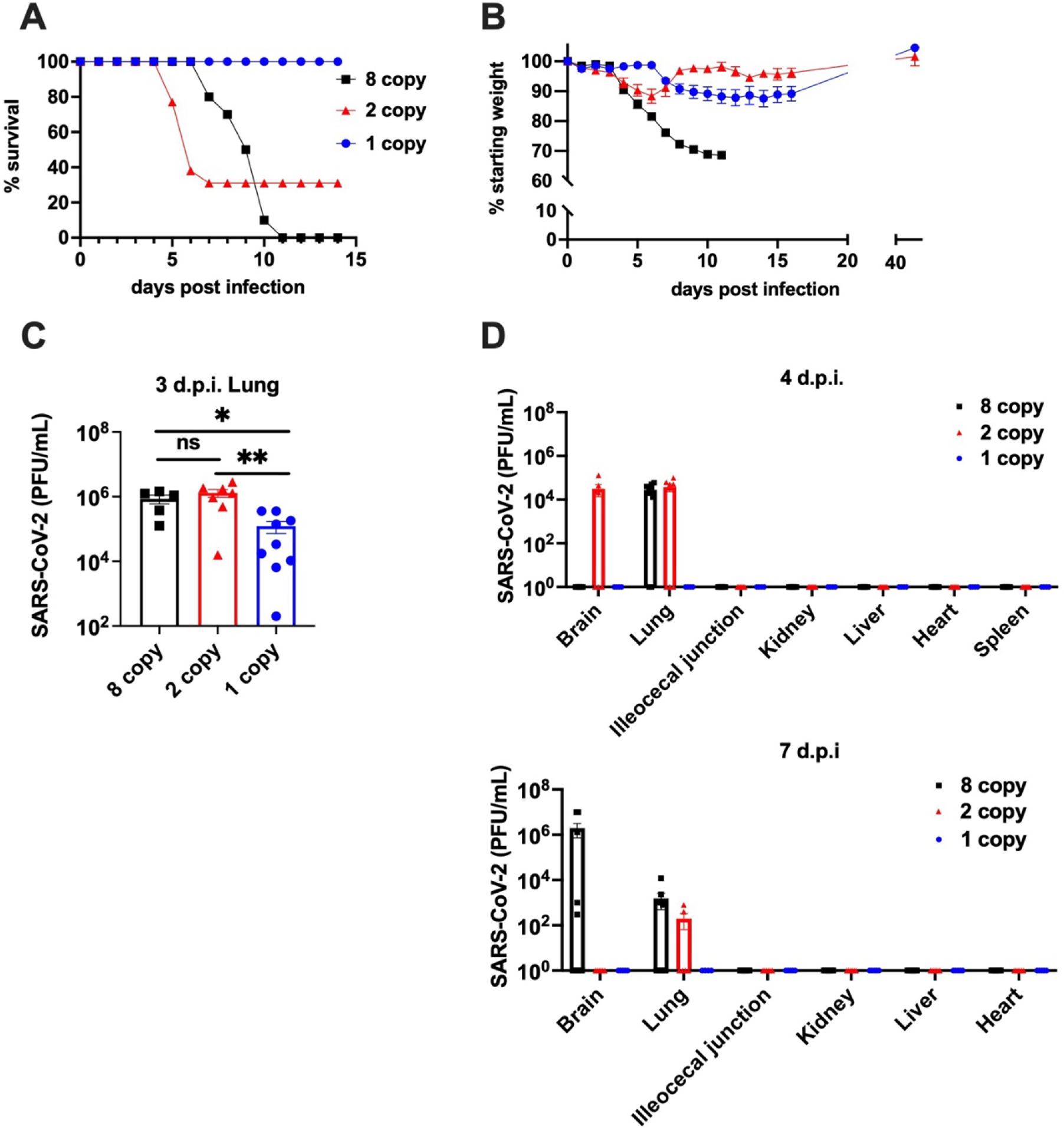
SARS-CoV-2 pathogenesis depends on the hACE2 copy number in transgenic mice. Male and female mice of each transgenic mouse line were intranasally infected with 10^3^ PFU of SARS-CoV-2 and monitored daily for (A) survival and (B) weight loss (1-hACE2-Tg n=10; 2-hACE2-Tg n=13; 8-hACE2-Tg n=10). Indicated tissues were harvested from infected mice on (C) 3 d.p.i., (D, top) 4 d.p.i, and (D, bottom) 7 d.p.i. and viral burden was quantified by plaque assay. Results are mean ± SEM for 4-11 mice per group. A Mann-Whitney test was performed to compare groups * p<0.05; ** p<0.008.

Previous studies have shown that the brain is a major site of viral replication in 8-hACE2-Tg mice^25^. To determine if transgene copy number impacts SARS-CoV-2 tissue tropism and viral spread, we measured viral loads in various organs known to express hACE2 in this model^4^ (**Figure 2 C, D**). Lung viral titers on day 3 post infection were comparable between 8-hACE2-Tg and 2-hACE2-Tg mice but were significantly lower in 1-hACE2-Tg mice (**Fig. 2C**). Infectious virus was below our limit of detection on day 4 p.i. in the ileocecum junction, kidney, liver, spleen, or heart. On 4 d.p.i. virus was detected in the brain of 2-hACE2-Tg mice but not in the brains of 8-hACE2-Tg or 1-hACE2-Tg mice **Fig. 2D**. However, by 7 d.p.i. 8-hACE2-Tg mice developed high viral brain titers, whereas the infection in the brains of surviving 2-hACE2-Tg mice had resolved **(Fig. 2D)**. Virus was not detectable in the brain of 1-hACE2-Tg mice at either of the time points analyzed (**Fig. 2D)**. The absence of viral replication in the brain of 1-hACE2-Tg mice, alongside of their survival despite significant lung titers on day 3, is consistent with the idea that brain infection determines the excess mortality in a transgene-dependent manner.

### Primary and memory responses in low-copy number hACE2-Tg mice

The commercially available 8-hACE2-Tg mice were widely used for vaccine and therapeutics testing^5,6,7,8,9^. However, due to their rapid and high mortality, they are not a good model for evaluating the adaptive immune response. We investigated the ability of the low copy number mice to mount primary and memory immune responses against SARS-CoV-2. We decided to focus our evaluation of immune responses to the 1-hACE2-Tg mice due to their 100% survival rate. Of note, we were also able to measure both T and B cell responses in 8- and 2-hACE2-Tg mice infected with the Omicron variant which is less lethal in mice **(Supplemental figure 1)**. For primary cellular immune responses against SARS-CoV-2 USA-WA1/2020, we found detectable SARS-CoV-2 spike-specific CD8+ T cells in peripheral blood on d7 post SARS-CoV-2 infection, as measured by H-2K^b^: VNFNFNGL/Spike_539-546_ tetramer binding^26^. Such responses ranged between 0.1 and 0.9% of blood CD8+ cells (**Fig. 3A**). 3 months post infection, we evaluated cellular and humoral memory immune responses. Splenocytes were stimulated with a SARS-CoV-2 spike peptide pool and CD4 and CD8 T cell production of IFNγ and TNFα was quantified. 1-hACE2-Tg mice exhibited a high frequency of spike-specific memory T cells secreting IFNγ and TNFα upon antigen stimulation **(Figure 3B)**. Finally, 1-hACE2-Tg mice also exhibited detectable neutralizing titers against SARS-CoV-2 and significantly higher levels of spike receptor binding domain (RBD)-specific IgG compared to mock infected mice measured at a memory time point, 3 months p.i. **(Figure 3C, D)**. We conclude that 1-hACE2-Tg mice produce detectable primary and memory immune responses upon intranasal infection with SARS-CoV-2.

**Figure 3.**
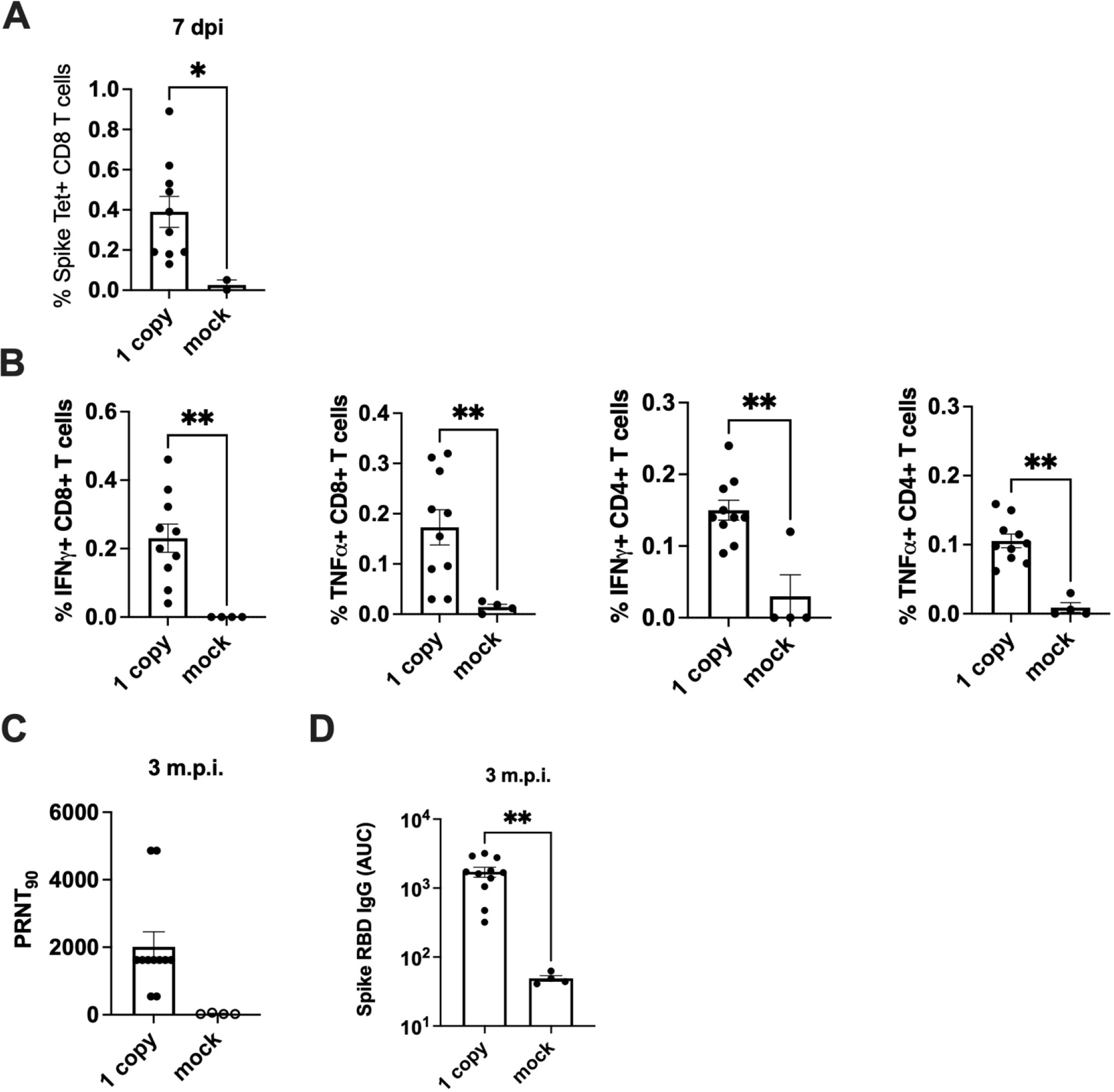
Primary and memory responses in 1-hACE2-Tg mice. On day 7 post infection, splenocytes were harvested and spike-tetramer specific CD8+ T cells in the spleen were quantified using flow cytometry (A). 3 months post infection, splenocytes and serum were collected from SARS-CoV-2 infected and mock infected 1-hACE2-Tg mice (B-D, infected n=11, mock infected n=4). (B) CD8+ and CD4+ T cell production of IFNγ and TNFα was measured upon stimulation with a USA-WA1/2020 spike peptide pool. (C) SARS-CoV-2 plaque reduction neutralization assays and (D) anti-spike RBD IgG ELISAs were performed on serum. All data presented are mean ± SEM. A Mann-Whitney test was performed to compare groups * p<0.05; ** p<0.008.

## Discussion

In this report, we detail generation and characterization of low copy-number hACE2-Tg mice as a model for infection with coronaviruses that use hACE2 as its cellular entry receptor, as assessed in the model of SARS-CoV-2 infection. We found that gene dosage of hACE2 directly and sharply influenced SARS-CoV-2 susceptibility, as well as other features of disease pathogenesis. Thus, in comparison with the established and widely available 8-hACE2-Tg mice^4^, 2-hACE2-Tg and 1-hACE2-Tg mice exhibited gradually reduced mortality and, for 1-hACE2-Tg mice, reduced virus replication in the lungs. From our limited analysis with a single infection dose of SARS-CoV-2, it appeared likely that gene dosage may exhibit a threshold effect. We speculate that this may be so for two reasons. First, lung titers were high and similar in 8-hACE2-Tg and 2-hACE2-Tg mice, but lower in 1-hACE2-Tg mice. Second, the differences in mortality and brain replication were both more pronounced between the 2-hACE2-Tg and 1-hACE2-Tg lines of mice, compared to the differences between 8-hACE2-Tg and 2-hACE2-Tg mice.

Most strikingly, we could not detect any brain virus replication on either d4 or d7 p.i. in 1-hACE2-Tg mice. An interesting, and at the surface paradoxical, difference was seen between 2-hACE2-Tg and 8-hACE2-Tg mice with regard to brain virus replication, where the virus was detected earlier (d4), but at lower levels, in the brains of 2-hACE2-Tg mice, compared to the 8-hACE2-Tg copy mice, who exhibited no detectable virus on d4 but high levels on d7. We hypothesize that this delay in brain infection in 8-hACE2-Tg mice may be due to the abundant availability of hACE2 in the lungs and respiratory tissues of these mice, leading to a delay of virus egress from the respiratory tissues and its late seeding of the CNS because the newly produced and released virus would have abundant binding targets in the respiratory environment. Conversely, a lower abundance of hACE2 receptors for virus in 2-hACE2-Tg would be more conducive to an earlier virus egress and dissemination to the brain. Of interest, we could not recover infectious virus from any organs other than the lung and the brain, in any of the transgenic mouse lines examined. Additional experiments varying the infectious dose will be required to further substantiate the above hypotheses and speculations.

Low-copy hACE2-transgenic mice were also able to generate detectable immune responses against SARS-CoV-2 in response to SARS-CoV-2 infection, and, in preliminary experiments, vaccination (data not shown). Specifically, primary CD8 T cell responses were readily detectable on day 7 following infection as judged by percentages of CD8 T cells staining with the H-2K^b^: VNFNFNGL/Spike^539-546^ tetramer. Such responses exhibited decreasing abundance with the reduction of transgene copy numbers yet were significantly higher over the background found in uninfected mice. We further detected memory immune responses in 1-hACE2-Tg mice, where memory CD8 responses were lower, but nonetheless clearly distinguishable from the background. We found similarly detectable memory CD4 and anti-S RBD antibody responses in these animals 3 months post infection.

Based on all the above, we conclude that the low copy hACE2-Tg mice, and in particular the 1-hACE2-Tg line, represent an optimized model for studies of pathogenesis and immunity against infection with viruses that use this molecule as their primary cellular entry receptors.

## Materials and Methods

### Mice

The transgenic vector, courtesy of Stanley Perlman and Paul B. McCray, Jr.^4^, containing the human hACE2 cDNA under the regulation of the human cytokeratin 18 (K18) promoter was injected into C57BL6/NJ mouse embryos using pronucleus microinjection. Fertilized eggs were collected from the oviducts of super-ovulated C57BL6/NJ females. Microinjections were performed by continuous flow injection of 1ng/µL exogenous DNA into the pronucleus of 1-cell zygotes. Genomic DNA was purified from tail tissue of newborn mice then PCR was performed to confirm transgene integration using the forward primer ACCTGGCTGAAAGACCAGAACAAG and reverse primer AATTAGCCACTCGCACATCC. Mouse strains were propagated of transgene-positive offspring by mating with C57BL6/NJ mice, maintaining heterozygosity of the offspring.

### Determination of *hACE2* copy number

Genomic DNA was isolated from tail tissue from each founder line of transgenic mice, C57BL6 mice, and K18-hACE2 mice purchased from Jackson Laboratory and was used as a template for TaqMan quantitative PCR. The primers and probe specific for *hACE2* (Life Technologies) are as follows: forward primer, TCCTAACCAGCCCCCTGTT; reverse primer, TGACAATGCCAACCACTATCACT; probe, ATATGGCTGATTGTTTTTGGAGTTGTGATGGG.

A single-copy reference gene, mouse Tfrc, was also quantified using the same templates and on the same reaction plate as *hACE2*. Each DNA sample was reacted in quadruplicate with a Fluorescein amidite (FAM) labeled transgene probe and a VIC labeled single-copy reference gene probe. A rodent glyceraldehyde-3-phosphate dehydrogenase (GAPDH) primer and probe set (Applied Biosystems) were included in each reaction to normalize the DNA added. ABI CopyCaller software was used to calculate the delta-delta-Ct values which were used to determine the transgene copy number.

### Extraction of total RNA and quantitative reverse transcription-PCR

Lung and brain tissues (5mm) from C57BL6 (wild type), F32 line (2 hACE2 copies), M4 line (1 hACE2 copy), and K18-hACE2 mice purchased from Jackson Laboratory (8 hACE2 copies) were snap-frozen on dry ice. The frozen tissues were submitted to the University of Arizona Genomics Core (UAGC) for RNA extraction and Quality Control using the Agilent Bioanalyzer (RNA 6000 Nano chips). After confirming RIN scores of 7.0+ for all samples, cDNA templates were generated using the Invitrogen SuperScript IV First Strand Synthesis kit. cDNA was pooled in triplicate with ABI-designed Taqman gene expression assays. Three assays were carried out on each well: GAPDH as an endogenous control, human- and mouse-specific ACE2 as the experimental genes. The human ACE2 was synthesized with the NED fluorophore allowing all three genes to be run as a triplicate. Forward primers were 5’-TCCTAACCAGCCCCCTGTT-3’ for hACE2 and 5’-TCTGCCACCCCACAGCTT-3’ for mACE2; reverse primers were 5’-TGACAATGCCAACCACTATCACT-3’ for hACE2 and 5’-GGCTGTCAAGAAGTTGTCCATTG-3’ for mACE2; probes were 5’-ATATGGCTGATTGTTTTTGGAGTTGTGATGGG-3’ for hACE2 and 5’-CACGGAGACTTCAGAATCAAGATC-3’ for mACE2. Delta-Delta Ct analysis was used to determine human vs mouse ACE2 expression. The differential is expressed as 2^(level of hACE2 – level of mouse ACE2)^ both tested against the ACTB reference gene.

### Infection and animal monitoring

The SARS-CoV-2 USA-WA1/2020 isolate was deposited by Dr. Natalie J. Thornburg at the Centers for Disease Control and prevention and obtained from the World Reference Center for Emerging Viruses and Arboviruses (WRCEVA) at The University of Texas Medical Branch. The virus was propagated and titered on Vero CCL-81 cells. Mice of both sexes, 8-12 weeks of age, were anesthetized with ketamine/xylazine and infected intranasally with 10^3^ plaque forming units (PFU) SARS-CoV-2 in 50 μL saline or diluent alone (mock infected). Mice were monitored and weighed daily. All experiments with SARS-CoV-2 were performed in either a Biosafety Level 3 or Animal Biosafety Level 3 laboratory at the University of Arizona.

### Viral plaque assay

For USA-WA-1/2020 and BA.1 (B.1.1.529.1 Omicron) titration, Vero cells (ATCC # CCL-81) were plated in 12 well plates at a density of 2×10^6^ cells per plate and incubated overnight at 37 °C in 5% CO_2._ Tissue homogenate supernatants were serially diluted in media (Dulbecco’s Modified Eagle Medium (DMEM) supplemented with 5% fetal bovine serum (FBS) and 1μM sodium pyruvate) then added to cells and incubated at 37 °C with 5% CO_2_ for 1 hour. Plates were overlayed with 1% methylcellulose supplemented with 1μM sodium pyruvate, 5% fetal bovine serum, penicillin, and streptomycin and incubated at 37ºC with 5% CO_2_ for 72 hours. Methylcellulose overlays were removed, plates were fixed in Neutral Buffered Formalin for 30 minutes, and plaques were visualized by staining with 0.9% (w/v) crystal violet in ethanol. Viral titers were quantified as PFU per mL of tissue.

### Plaque Reduction Neutralization Titer assay

Vero cells (ATCC # CCL-81) were plated in 96 well tissue culture plates at a density of 2×10^6^ cells per plate and incubated overnight at 37 °C in 5% CO_2._ Serial dilutions of mouse serum samples were incubated with 100 PFU of SARS-CoV-2 for 1 h at 37 °C. Serum dilutions/virus mixtures were transferred to confluent Vero cell monolayers and incubated for 2 h at 37 °C with 5% CO_2._ Plates were overlayed with 1% methylcellulose supplemented with 1μM sodium pyruvate, 5% FBS, penicillin, and streptomycin then incubated at 37ºC with 5% CO_2_ for 72 hours. After 72 hours, plates were fixed with 10% Neutral Buffered Formalin for 30 min and plaques were visualized by staining with 0.9% (w/v) crystal violet in ethanol.

Plates were imaged using an ImmunoSpot Versa (Cellular Technology Limited, Cleveland, OH) plate reader. Results are expressed as the serum dilution that produced a 90% reduction of plaques.

### ELISA

SARS-CoV-2 Spike RBD protein (GenScript #Z03479) was immobilized on high-absorbency 96-well plates at 5 ng/mL and incubated at 4°C overnight. Plates were blocked with 1% non-fat dehydrated milk in sterile PBS for 1 hour at room temperature, washed with PBS containing 0.05% Tween-20, and overlaid with serial dilutions of serum for 1 hour at room temperature. Next, plates were washed thoroughly and incubated with an anti-mouse IgG (H+L) peroxidase-labeled antibody (SeraCare #5450-0011) at a concentration of 1:1000 for 1 hour at room temperature. Plates were washed with PBS-Tween solution followed by PBS wash and developed using 3,3’, 5,5’-tetramethylbenzidine solution prior to quenching with 1M H_2_SO_4_. Plates were then read for 450nm absorbance. GraphPad Prism software was used to calculate the Area Under the Curve values using a technical baseline of 0.05 based on OD readings from control wells without serum.

### T cell peptide stimulation and Flow Cytometry detection of intracellular cytokine production and antigen-specific cells

Blood was collected retro-orbitally from mice. Hypotonic lysis was performed to lyse red blood cells. Cells were resuspended and stained overnight at 4°C with antibodies against CD3, CD4, CD8 and H-2K^b^: VNFNFNGL/Spike_539-546_ tetramer (NIH Tetramer Core Facility, Emory University, Atlanta, GA) followed by Live/Dead viability dye and fixation^26^. Spleens were collected and disassociated using 40μM mesh cell strainers. Single cell suspensions from spleens were split and stimulated with a SARS-CoV-2 Spike pool or just media alone (no stimulation condition) in the presence of 2-Mercaptoethanol (BME) and protein transport inhibitor for 6 hours at 37°C. Cells were washed and stained with antibodies against CD3, CD4, and CD8 followed by Live/Dead viability dye. Cells were then subjected to fixation and permeabilization for intracellular staining with antibodies against IFNγ, and TNFα. Samples were acquired on a BD LSR Fortessa cytometer (BD Biosciences) and data was analyzed using FlowJo software.

## ABBREVIATIONS

8-, 2- and 1-hACE2-Tg: mice carrying 8, 2 and 1 hACE2 transgene copies, respectively
hACE-2: human angiotensin converting enzyme 2
i.n.: intranasal
p.f.u.: plaque-forming units
p.i.: post infection
d.p.i.: days post infection
Tg: transgenic.

---

**Supplemental figure 1.**
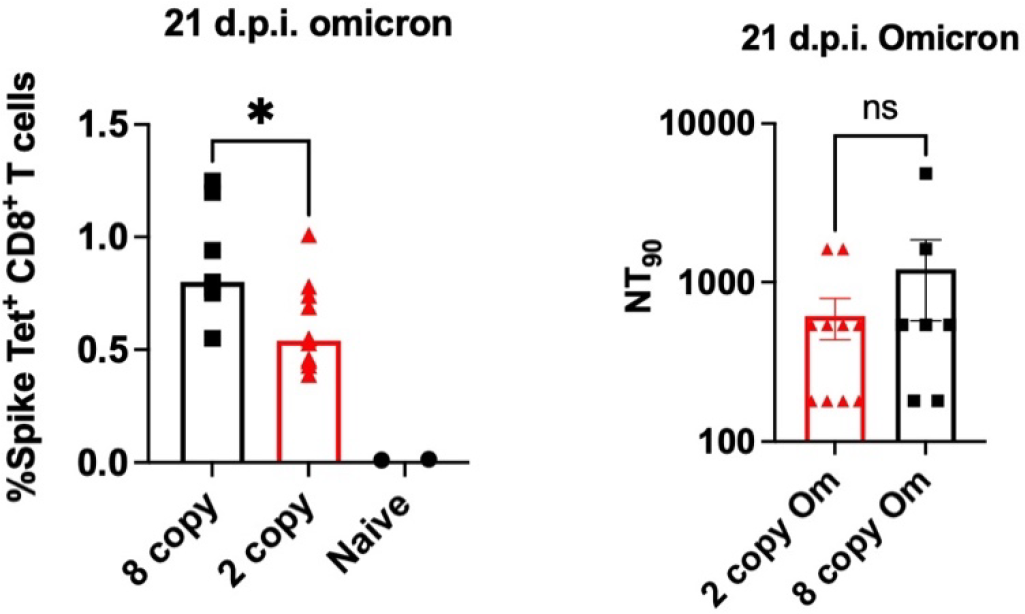
Omicron infection in low-copy number hACE2-Tg mice. 2- and 8-hACE2-Tg mice were intranasally infected with the SARS-CoV-2 omicron variant. On day 21 post infection, spike-tetramer specific T cells in the spleen and serum neutralizing titers were quantified. All data presented are mean ± SEM. A Mann-Whitney test was performed to compare groups * p<0.05.

